# Microfluidic Separation of Adipocytes

**DOI:** 10.64898/2026.03.07.710291

**Authors:** Jason P. Beech, Mathis Neuhaus, Karin G. Stenkula, Jonas O. Tegenfeldt

## Abstract

Adipocyte size is an independent predictor of several metabolic disorders, including type 2 diabetes, liver and cardiovascular diseases. However, technical limitations due to the fragile nature of mature adipocytes have restricted the functional analyses of size-separated adipocytes using conventional methods. Therefore, we have developed a microfluidic device, based on deterministic lateral displacement, for sorting intact, mature adipocytes. Cell-size distribution was determined from time-lapse recordings inside the device, in separate outlets, and by Coulter counter analysis of the collected cell fractions. This approach allowed size-separation with minimal size-overlap with mean diameters of (small fraction) 47 µm and (large fraction) 82 µm based on Coulter counter measurements. Viability of the separated cells was verified by insulin stimulation and western blotting of key insulin signaling proteins. The sample recovery, comparing input versus output material, was relatively high, 42% for the large fraction with a purity of 93%.

We demonstrate that microfluidics is a suitable approach to overcome the limitations of sorting mature adipocytes according to size. Together, the high recovery rate, high throughput capacity, accurate separation and the fact that the cells maintained hormonal response after sorting provides compelling evidence of the strength and usability of the microfluidic approach for exploring adipocyte function in relation to size.

## 2. Introduction

Obesity and its associated metabolic complications represent some of the most significant and growing public health threats. Current estimates suggest that approximately 40% of diabetes cases are directly linked to obesity (1). Adipocyte dysfunction is a fundamental defect characterizing these diseases (2). In adaptation to energy surplus, adipose tissue expands both by increasing the number of adipocytes (hyperplasia) and adipocyte size (hypertrophy) (3). In humans, adipocytes with sizes ranging from less than 20 µm to larger than 300 µm have been observed, which extrapolates to a several thousand-fold difference in cell volume (4). In periods of caloric restriction, adipose tissue mass usually adapts by decreasing adipocyte size rather than cell number (3). Adipocyte size is an independent predictor of a plethora of medical complications comprising type 2 diabetes, hepatic complications, metabolic syndrome, and cardiovascular diseases (5-7). Large, hypertrophic adipocytes are commonly believed to be less insulin sensitive (8), have higher basal and blunted catecholamine-stimulated lipolysis (9), display a profile of increased pro-inflammatory adipokine and cytokine secretion (10), and are characterized by limited expandability. In contrast, small adipocytes are generally considered more insulin-responsive and display increased secretion of favorable adipokines (8, 10). Due to these size-driven discrepancies in adipocyte characteristics, decades of research have sought to separate adipocytes by size. However, the fragile nature of fat cells makes common separation approaches like FACS or modern single-cell analysis platforms cumbersome. In 1970, Björntorp and Karlsson were amongst the first to separate large and small cells from the same fat tissue sample in dialysis tubing based on differences in buoyancy (11). Over the years, several other studies have used separation funnels or nylon meshes of different pore sizes to collect pools of small and large adipocytes (12). Frequent problems with all these approaches are cell breakage, cell aggregation (small cells sticking to large), cell pliability, and the large required amounts of input material compared to the limited output.

In the current study, we aimed to overcome these technical limitations using microfluidics. Microfluidic technologies have become widely used for size-based cell separation due to their ability to precisely control fluid flow and particle trajectories at the microscale (13-15). By leveraging mechanisms such as deterministic lateral displacement (DLD) (16, 17), inertial focusing (18, 19), and microfiltration (20), these platforms enable label-free and continuous sorting of heterogeneous cell populations. Such approaches have been successfully applied to applications including blood fractionation and isolation of rare cells, offering advantages over conventional methods in terms of throughput, sample handling, and system integration (21). Although microfluidic technologies have been widely applied to size-based separation of many cell types, their use for the separation of adipocytes remains limited. Most published work involving adipose tissue has focused on microfluidic culture platforms and organ-on-chip models, rather than separation. For example, Li and Easley reviewed emerging microfluidic systems for studying adipocyte function and adipose tissue dynamics, emphasizing the technical challenges associated with handling mature adipocytes in confined microchannels (22). Similarly, Yang *et al*. (23) and Compera *et al*. (24) developed microfluidic platforms for three-dimensional adipose tissue culture and on-chip adipogenesis, while Huff *et al*. (25) demonstrated perfused fat-on-a-chip systems using primary mature adipocytes. By contrast, studies specifically addressing size-based microfluidic separation of mature adipocytes are largely absent from the literature. Existing separation approaches more commonly target adipose-derived stromal or stem cell populations using inertial or hydrodynamic techniques, rather than isolating buoyant, fragile mature adipocytes themselves. Consequently, despite the maturity of size-based microfluidic sorting for other cell types, its application to adipocytes remains underexplored.

Here, we demonstrate that microfluidic technologies, specifically Deterministic Lateral Displacement (DLD), could provide a viable strategy to overcome the longstanding challenges of adipocyte sorting, thereby establishing a foundation for future investigations that may yield novel insights into adipocyte biology.

## 3. Results and Discussion

We demonstrate that microfluidics can be a suitable approach to separate viable adipocytes by their cellular size. The device demonstrated stable flow, producing reproducible flow profiles and consistent cell trajectories within the DLD array (Figure 1 and SI movies 1 and 2). Handling buoyant, deformable adipocytes in DLD devices represents a significant technical challenge compared with previously studied cell types. Because mature adipocytes rapidly separate from the carrier buffer due to buoyancy, continuous gentle mixing of the sample reservoir using a magnetic stirrer was required to ensure uniform cell entry into the device. Under these operating conditions, adipocytes passed through the array without detectable fouling or accumulation on channel walls or pillar structures, indicating cell compatibility with device surface properties and low damaging shear stresses (see SI movie 2). Although occasional lipid droplets were detected, their low abundance and distribution suggest they primarily originated from upstream handling rather than device-induced rupture. These observations indicate that the platform enables stable processing of fragile, buoyant adipocytes while preserving conditions necessary for size-dependent deterministic displacement.

**Figure 1:**
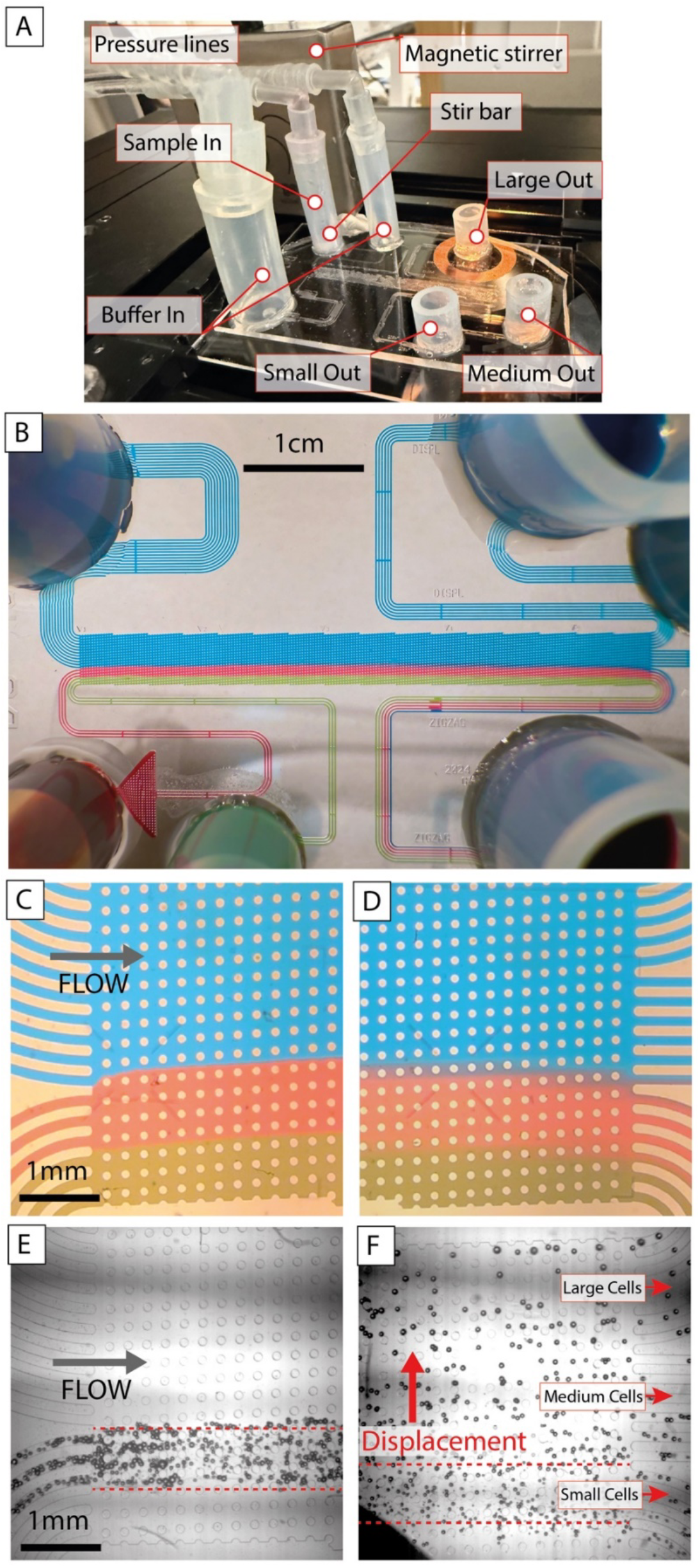
Overview of fluidics device with examples of separation results. (A) Photo of actual device showing inlets, outlets and tubes for pressure control. A small magnetic stir bar is used in the sample inlet reservoir to stop the fat cells separating from the buffer due to the large difference in bouyancy. (B) Food colouring is used to highlight the laminar flow of the carrying fluid from the three inlets and to show how it exits the device. A small amount of diffusion can be observed. (C-D) Close-ups of flow. Note the pillar array that gives rise to the DLD mechanism. (E) Cells entering separation array from the left and flow through the array to the right. (F) Cells exiting the separation array after size-based displacement. Larger cells are displaced towards their left (upwards in the image).

Our automated analysis of images of 35,000 cells in the three different device outlets (see SI movie 3) shows a distinct separation of the measured particle sizes (Figure 2A-B) where we show small (blue) and large (green) fractions, The intermediate (Guard) fraction is shown in the routing probability analysis in the SI. The overlap between the small and large fraction particle sizes is minor. The intermediate outlet was intentionally designed to collect particles near the critical diameter (*D*_*C*_), reducing cross-contamination and improving the purity of the small and large fractions and functions well in this respect.

**Figure 2:**
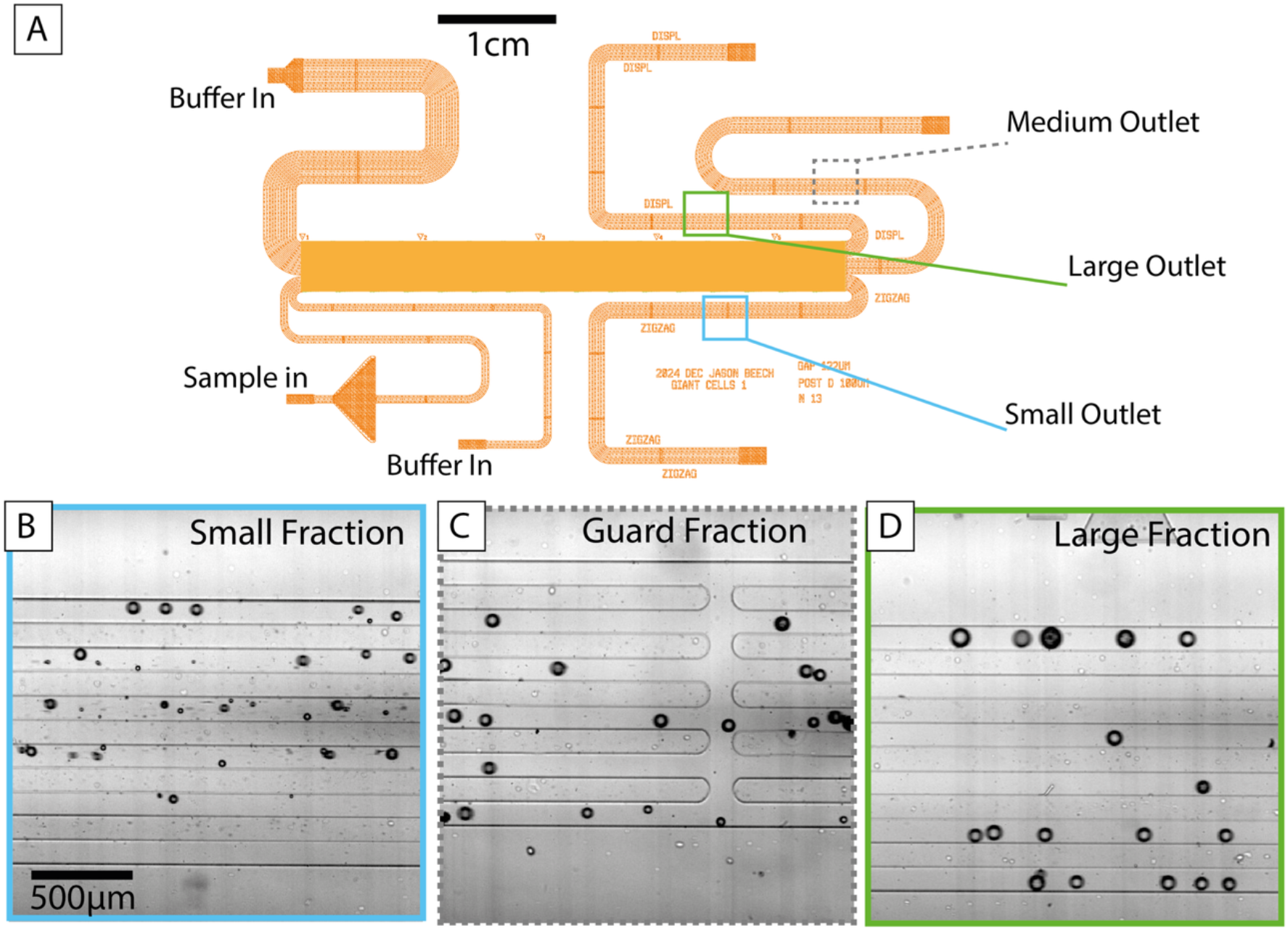
How images are acquired in the three device outlets. (A) An overview of the device. Colored squares show where images are recorded. (B-C) Representative frames from images acquired in the three outlets. These images are used for size distribution measurements. Actual cell counts (35,000 in total).

Adipocytes accumulating in the outlets (small, large, and intermediate) were collected and used for subsequent Coulter counter analysis. Here, we could again demonstrate a clear separation in the measured particle size from small and large fractions (Figure 3A-C). Using two high-throughput cell-size analysis tools, we could independently confirm the robust and consistent separation of the fractions. The small fraction had a mean diameter of 56µm, the intermediate fraction of 67µm, and the large fraction of 76µm, which extrapolates to an approximately 2.5-fold difference in mean cellular volume comparing the small and the large fractions (Figure 3C). This is comparable to separations previously obtained (10). Yet, we believe that this approach, with further modifications, will allow for a clearer separation of the fractions, higher scalability, and potential automation. From previous experience with the floatation-based method, used for example by Skurk et al. (10) and Jernås et al. (12), we found that a major limitation is the high amount of input material required and the limited number of cells obtained in the small-cell fraction. Here, we used approximately 500µL of packed adipocytes and sequentially ran those cells through our microfluidics device at a pressure of 20mbar on each of the inlets for 1h, yielding approximately 150µL packed adipocytes in the small fraction and 100µL in the large fraction. This ratio of input to output is superior compared to previous approaches using 5-40g of adipose tissue for cell size separation (8, 10, 12).

**Figure 3:**
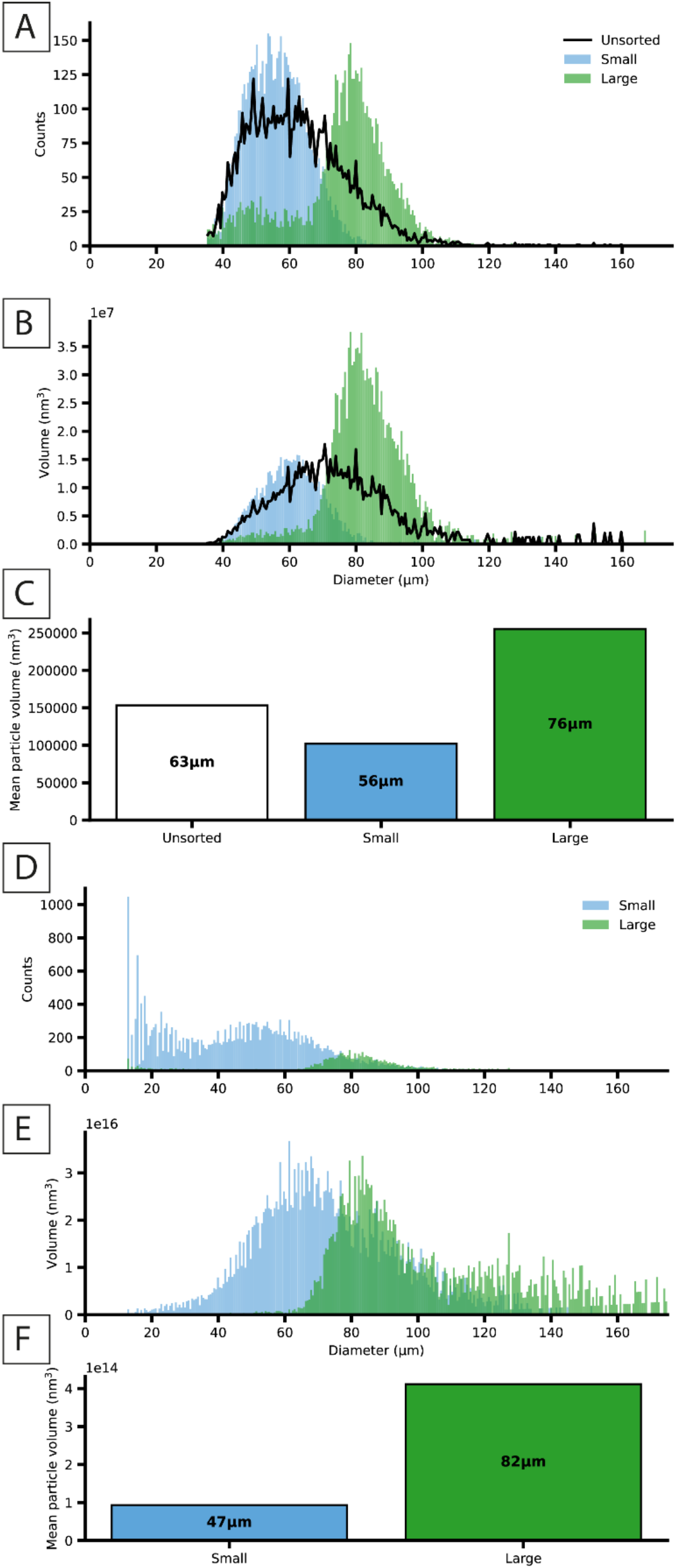
Measured fat cell diameter distributions using a Coulter counter and in separation device. (A) Size distribution of 6000 counts of osmium-fixated adipocytes before (black line) and after separation. Small fraction in blue and large in green. All bins are 0.55µm (B) Volume of particles measured per bin in 6000 counts. (C) Mean particle volume before and in the small and large adipocyte fractions, in the middle of each bar is the mean particle diameter in micrometers. The middle (guard) fraction is not considered here as it is designed to increase the purity of the other two fractions. (D) Diameters of unfixed adipocytes measured directly after sorting (in outlet channels), approx. 35 000 cell counts. (E) Volume calculated from diameters assuming perfect spheres. Mean cell volumes for small and large fractions with mean cell volume displayed in the center of the bars.

Although the device was designed with a theoretical critical diameter (D_C_) of 50 µm, the experimentally observed separation threshold was closer to ∼70 µm based on both Coulter counter measurements and routing probabilities; a statistical analysis of the probabilities of cells of a given size to exit via either of the three outlets described in the SI. Several factors likely contribute to this discrepancy. First, Coulter counter measurements of osmium-fixed adipocytes are known to overestimate cell diameter due to fixation-induced swelling and the conductive shell formed during osmium treatment (32). Second, diameter estimation from bright-field image segmentation may introduce systematic overestimation arising from optical blur, thresholding, and the difficulty of defining precise cell boundaries. Finally, adipocytes are highly deformable and can adopt non-spherical shapes while traversing the DLD array, resulting in an effective hydrodynamic diameter smaller than their apparent static diameter(33), although appreciable deformations were not observed (see below for further discussions on deformability). Together, these effects would shift the apparent cutoff towards larger measured diameters compared with the theoretical *D*_*C*_.

Routing probability analysis allows us to calculate yields and purities for the different fractions (see SI). We get purities of 97 and 93% (small, large) and yields of 66 and 42%. These numbers highlight the increased purity enabled by the inclusion of the guard fraction but also the cost in reduced yield since most “lost” cells are collected there.

The mean flow velocity in our device is approximately 1 mm s^−1^, corresponding to a Reynolds number of ∼0.1 for a characteristic gap size of 122 µm, indicating laminar creeping flow. Assuming a parabolic velocity profile, this corresponds to an estimated wall shear rate of ∼40 s^−1^ and shear stress of ∼0.04 Pa (assuming water-like viscosity). These stresses are several orders of magnitude smaller than those applied in microfluidic deformability assays such as real-time deformability cytometry (RT-DC), where cells experience hydrodynamic stresses of ∼1–100 Pa (34, 35) Given that adipocyte-like cells exhibit kPa-scale stiffness (36), shear-induced deformation is therefore expected to be negligible. The low Reynolds number and modest shear rates further suggest substantial scope to increase throughput by raising flow velocities while remaining in the laminar regime required for DLD and well below stresses typically used to mechanically deform cells.

To investigate the viability of adipocytes after separation, we measured their response to insulin using western blotting of key insulin signaling targets (Figure 4A-C). Total protein staining reveals that the amount of protein (disregarding BSA contamination from lysate preparation) is lower in the small than in the “before-sorting” fraction and even lower in the large fraction (Figure 4A), while BSA contamination is higher in the fractions after separation. Caveolin 1 and Fatty Acid Synthase (FAS) were used as markers of adipocytes, and serine 473 protein kinase B phosphorylation (PKB S473) and Akt substrate of 160 kDa threonine 642 phosphorylation (AS160 Thr642) as measures of insulin responsiveness (Figure 4B-C). We observed a clear induction of PKB and AS160 phosphorylation in all fractions, before and after separation, upon insulin-stimulation. These results demonstrate that the obtained adipocyte fractions respond to insulin and are viable.

**Figure 4:**
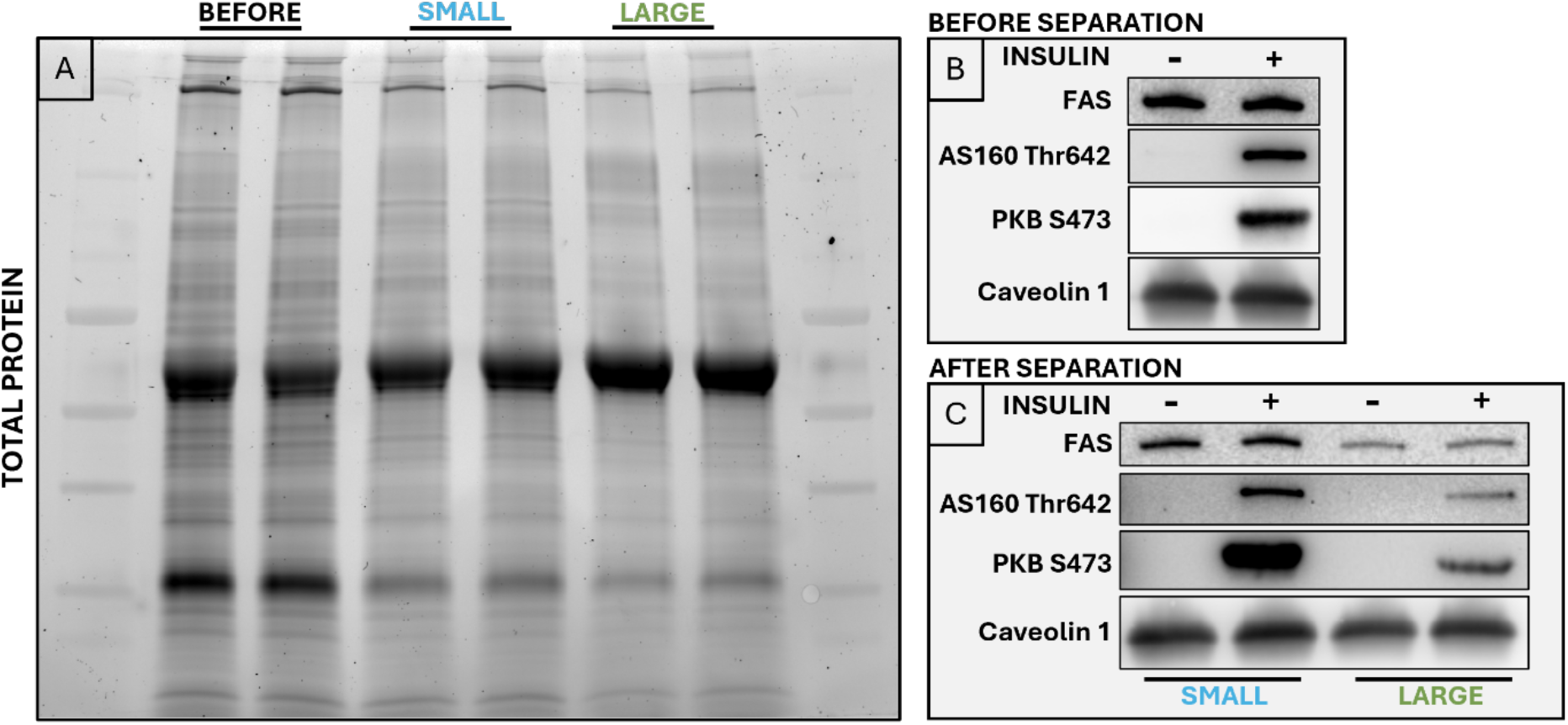
Western Blotting of small and large adipocytes. (A) Total protein stain of adipocyte lysates, the cell fractions: before, small, and large. (B) Western Blotting before separation in basal and insulin-stimulated (10nM, 5min) adipocyte lysates. C) Western Blotting after separation in basal and insulin-stimulated (10nM, 5min) small and large adipocyte lysates. Note that images before (B) and after (C) separation were taken at different exposures and should thus not be directly compared.

In conclusion, we demonstrate that microfluidics is a suitable approach to overcome the current limitations arising when trying to separate viable adipocytes by their size. The fact that the adipocytes maintained their integrity and hormonal response after sorting emphasizes the strength and usability of microfluidics for exploring adipocyte function in relation to size.

While the present approach was shown to be efficient for size-sorting, some limitations remain, which could be resolved with further optimization. The minor size overlap in the small and large fractions could potentially be reduced further by fine-tuning the *D*_*C*_ range of the device. Changing the ratios between small, guard and large fraction outlet geometries would also allow for the optimization of purity or yield, depending on the requirements thereof. As discussed above there is room for increasing throughput also by increasing flow rates. A limitation in the current study was the size of the inlet and outlet reservoirs, which were designed to allow sub-ml of sample (0.325mL) and required refilling during experiments, but this could easily be increased. Parallelization of fluidics devices, together with increased reservoir size and volume flow rates, provides a route to faster separations of larger volumes of cells.

## 4. Materials and Methods

### 4.1 Animals and ethics statement

Male C57BL/6J (Taconic, Ry, Denmark) were housed on a 12h light cycle with nonrestricted food and water supply. Mice were fed a chow-diet SAFE®A30 (SAFE® Complete Care Competence). Tissue collection was performed in the fed-state. At termination (12 weeks of age), inguinal and epididymal adipose tissue was excised and used for analysis. All animal procedures were approved by the Malmö/Lund Committee for Animal Experiment Ethics, Lund, Sweden (ethical permit 17479-24)

### 4.2 Adipocyte isolation

Primary adipocytes were isolated from epididymal and inguinal adipose tissue depots in Krebs Ringer Bicarbonate HEPES (KRBH) buffer, pH 7.4, containing 200nM adenosine, 5% (w/v) bovine serum albumin (BSA), and 1mg/ml collagenase type I (Worthington Biochemical Corporation collagenase type I, LS004194) using an established protocol (26). Collagenase digestion was performed for one hour at 37 °C and 120 rpm in a shaking water bath (Grant Instruments). Isolated cells were filtered through a 400µm nylon-mesh and washed four times in KRBH buffer containing 200nM adenosine and 5% (w/v) BSA. The isolated adipocytes from pooled epididymal and inguinal adipose tissue samples were used for all subsequent experiments.

### 4.3 Device design and fabrication

A photomask for UV lithography was fabricated by Delta Mask (Enschede, The Netherlands). The design was drawn using a combination of the Python library Nazca (27) and L-Edit v.18.3 (Tanner Research, Monrovia, California, USA). The DLD array was designed using **Error! Reference source not found**. (28) with G = 122µm and N=13 to have a D_C_ of 50µm. The array with edge modifications (29) was generated programmatically using Nazca and imported in L-Edit where the remainder of the design was added.

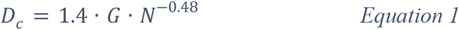

Inlets and outlets were designed to have matching fluidic resistances to ensure parallel, laminar flow into and out of the separation array. An intermediate outlet was incorporated to collect particles close to the critical diameter, acting effectively as a guard fraction as is well-known from standard liquid chromatography(30), improving separationof the extreme fractions. The resistances of the inlets/outlets were also designed to be higher than in the array itself since a theoretical analysis found this to provide more stability in the flow when tuning pressures (see SI).

A 4” silicon wafer was prepared by oxygen plasma treatment at 5 millibar for 60 s (Plasma Preen - North Brunswick, NJ, USA) and drying at 120 ºC. SU8 2015 was spun onto the wafer at 6000 rpm for 60 s, baked for 1 min at 65 ºC + 4 mins at 95 ºC, flood exposed for 20 s at 30 mW/cm2 (365 nm) and finally post exposure baked for 1 min at 65 ºC + 4 mins at 95 ºC. This layer is essential to ensure adhesion of the dry film resist that was subsequently applied to the wafer.

A 200 μm thick dry film resist (SUEX., K200, DJ Microlaminates, Sudbury, Massachusetts, USA) was laminated onto the wafer using a laminating machine (Catena 35, Acco UK Ltd, Buckinghamshire, UK) at 65 ºC. After a pre-exposure bake at 85 ºC on a hotplate (Model 1000-1 Precision Hot Plate, Electronic Micro Systems Ltd, West Midlands, UK) to relax the resist, exposure at 365 nm was performed in a contact mask aligner (Karl Suss MJB4 soft UV, Munich, Germany) for 40 s at a lamp power of 30 mW/cm2. Post exposure baking was done at 85 ºC for 12 minutes. In order to avoid mechanical stresses and delamination slow ramping of the temperature both up and down was done over at least 20 minutes. The SUEX film was development in mr-DEV 600 (Micro Resist Technology GmbH, Berlin, Germany) for 40 minutes plus 20 minutes in fresh developer followed by 5 mins in IPA (isopropylalcohol) and a final rinse in flowing IPA and drying with nitrogen. A final hard-bake was done on a hotplate by ramping from room temp to 95 ºC, leaving for 9 minutes and then allowing the wafer to cool slowly to room temperature. To reduce adhesion of PDMS (polydimethylsiloxane) to the master mould, an ALD (atomic layer deposition) system (Fiji – Plasma Enhanced ALD, Veeco, NY, USA) was used to deposit a layer of aluminium oxide (1 nm) followed by a monolayer of perfluorodecyltrichlorosilane (FDTS).

The microfluidic devices were fabricated by replica moulding of PDMS, (Sylgard 184, Dow Corning, Midland, MI, USA) at a mixture of 10:1, on the master mould. Curing was performed at 80 ºC for 60 minutes. Access holes were punched through the cast PDMS pieces and air plasma treatment (Zepto-BRS, Diener electronic GmbH & Co. KG, Ebhausen, Germany), of both PDMS (15 s) and glass slides (60 s) used to bond devices. Silicon tubes of sizes chosen to match volumetric flow rates were attached to the PDMS device.

### 4.4 Running Separations

Separation devices were first primed with running buffer (KRBH buffer containing 200nM adenosine and 5% (w/v) BSA). Cell suspensions were then introduced into the sample inlet using wide-bore pipette tips to minimize shear stress. To prevent adipocytes from creaming to the surface and thereby limiting their entry into the device, a small magnetic stir bar was placed in the sample reservoir and actuated using a magnetic stirrer positioned next to the reservoir to ensure continuous, gentle mixing. The buffer inlets hold 1.5mL (large) and 0.325mL (small), and the sample inlet holds 0.325mL. Buffer inlet and sample inlet were refilled multiple times during the run.

Flow through the device was generated by applying a pressure of 20 mbar to each inlet reservoir using an MFCS-4C pressure controller (Fluigent, Paris, France). Separated fractions were collected periodically from the outlet reservoirs using wide-bore pipette tips. During separation, real-time microscopy videos were acquired of cells traversing the microfluidic array to visualize sorting behavior, as well as within the outlet channels to enable subsequent size analysis.

All images were acquired with a Nikon epifluorescence microscope (Eclipse Ti microscope, Nikon Corporation, Tokyo, Japan) with a 4x objective (Nikon Plan Apo λ, NA 0.2), an EMCCD camera (iXon Life 897, Andor Technology, Belfast, Northern Ireland) with 512 x 512 pixels (16 µm x 16 µm) and sensor size 8.2mm x 8.2mm using the acquisition software NIS-elements AR. In order to minimize motion blur, images were captured at 2ms exposure times, and to ensure each image contained a unique set of particles (to ensure that no particles are counted twice in the size analysis), a delay of 100ms between frames was used.

### 4.5 Image Analysis

Particle segmentation was performed using a static background subtraction approach followed by adaptive thresholding and morphological cleanup to isolate individual particle objects in each frame. Object size and shape metrics were extracted from the resulting binary masks. Full details of the segmentation and measurement procedure are provided in the SI.

### 4.6 Insulin-stimulation and cell lysis

Fractions of isolated adipocytes before and after (small and large fraction) separation were briefly centrifuged at 200g to ensure packing and floatation of cells. Subsequently, cells were stimulated with 10nM insulin (NovoRapid®) for 5min at 37 °C, 120 rpm in a Grant Instruments water bath in KRBH buffer with 200nM adenosine but no added BSA. A fraction of cells was incubated in the same buffer without insulin and was considered “basal”. Basal and insulin-stimulated adipocytes were then lysed 1:1 (50µL packed cell volume) in a lysis buffer containing 50 mM Tris/HCl pH 7.5, 1 mM EGTA, 1 mM EDTA, 0.27 M sucrose, 1% NP-40, and complete protease inhibitor cocktail (Roche, Basel, Switzerland). Lysates were centrifuged for 10 min at 13,000g, and the infranatant was collected for further analysis. Bradford measurement was used to determine the total protein content.

### 4.7 Coulter counter measurement

Isolated adipocytes were fixed in 2% (w/v) osmium tetroxide to determine adipocyte size distribution using a Multisizer 4e Coulter Counter (Beckman-Coulter, Indianapolis, IN, USA) as described previously (31). After fixation, particles were washed (0.9% NaCl solution) through nylon-meshes with a pore size of 20µm and 250µm. Aliquots of adipocytes (about 40µL packed cell volume) before and after cell-size separation were subjected to Coulter counter analysis. Samples after separation comprise the small, intermediate, and large cell fractions. For each sample, the size of 6,000 particles was counted, from which we obtained the cell-size distribution curve. Data were analysed using linear bins (400 bins, bin size 0.55 μm), Multisizer3 version 3.53.

### 4.8 Western blot analysis

Equal amounts of protein lysates (15µg according to Bradford measurement) from the fractions: before, small, and large were loaded on stain-free TGX 4-15% gradient gels (No. 5678084, Bio-Rad, Hercules) in a sample buffer containing 1X Laemmli Sample Buffer (BioRad, #1610747) and 300mM 2-Mercaptoethanol (BioRad, #1610710). Separation of proteins was performed at 200V for 1h in Tris/Glycine/SDS Electrophoresis Buffer (BioRad, #1610772EDU). After separation, the stain-free gel was activated for 45s in a Bio-Rad ChemiDoc MP Imaging System, and proteins were transferred to an Immun-Blot® Low Fluorescence PVDF membrane (BioRad, #1620260). The membrane was blocked in 5% skim-milk powder for 1h at room temperature, washed in TBST-0.1% Tween20, and incubated in primary antibodies at 4 °C overnight. Primary antibodies against Fatty Acid Synthase (Cell Signaling Technology Cat# 3180, RRID:AB_2100796) diluted 1:2000 in TBST-5% BSA, Caveolin 1 (Cell Signaling Technology Cat# 3267, RRID:AB_2275453) 1:10,000, Phospho-Akt (Ser473) (Cell Signaling Technology Cat# 9271, RRID:AB_329825) 1:2000, and Phospho-AS160 (Thr642) (Cell Signaling Technology Cat# 4288, RRID:AB_10545274) 1:500 were used. Secondary antibody incubation was performed for 1h at room temperature with an anti-rabbit (No. 314060, Thermo Fisher Scientific, Waltham, MA, USA) horseradish peroxidase-conjugated secondary antibody diluted 1:2,500. After primary and secondary antibody incubation, membranes were washed 5x5min in TBST-0.1% Tween20. Enhanced chemiluminescence reagent (SuperSignal West Pico Chemiluminescent Substrates, Thermo Scientific) was used for developing the signals. Bio-Rad ChemiDoc MP Imaging System and Image Lab software (Bio-Rad Laboratories) were used for detection and quantification.

## Supporting information

Supplementary information regarding image segmentation and statistical analysis

## 5. Data availability statement

Numerical data for the Coulter counter experiments plus microscopy images used for all plots presented here are available in the following repository: https://doi.org/10.7910/DVN/M6DMVF

## 6. Code availability statement

The Python code used to segment and extract cell sizes from microscopy images, plus that to generate the routing probability analysis can be found in the following repository: https://doi.org/10.5281/zenodo.18789356

7. Acknowledgments

We would like to thank Maria Lindahl and Jieyu Zhang for their experimental support. We are also greatful to Bao Dang Ho for initiating this collaboration. This work was financially supported by the Swedish Research Council (2023-01779) and Strategic Research Areas EXODIAB (2009-1039) and NanoLund (staff01-2020, s01-2024), the Swedish Foundation for Strategic Research (IRC15-0067), The Royal Physiographic Society of Lund, and Albert Påhlsson Foundation. … and Crafoord for the equipment…

## 8. Author contributions

JPB, MN, KGS, JOT conceived and designed the study. JPB, MN, performed the experiments. JPB wrote python code and JPB, MN performed data analysis. Supervision, Resources, Project administration, KGS, JOT. All authors contributed in the writing of the final manuscript.

