## Supplementary information regarding image segmentation and statistical analysis for "Microfluidic Separation of Adipocytes"

### Microfluidic Separation of Adipocytes -Supporting Information-

#### Image segmentation and particle size extraction

Particle segmentation was performed on two-dimensional time-lapse TIFF stacks acquired at the outlets of the microfluidic device. Each stack consisted of grayscale images containing static microchannel structures and transient, dark, approximately spherical particles.

#### Background estimation and subtraction

To suppress static channel features while preserving moving particles, a static background image was estimated independently for each TIFF stack by computing a high-percentile projection across time (typically the 90th percentile at each pixel). Because particles appear darker than the background, this operation yields an estimate of the background intensity unaffected by transient particle passages. Each frame was then background-subtracted by computing the pixel-wise difference between the background image and the raw frame, followed by clipping of negative values to zero. This transformation rendered particles as positive-contrast objects on a near-zero background.

#### Noise suppression and thresholding

The background-subtracted frames were optionally smoothed using a spatial Gaussian filter ($\sigma \approx 1$ pixel) to reduce high-frequency noise without blurring particle boundaries. Segmentation was performed using **Otsu's method**, applied independently to each frame to compute an adaptive global threshold. Pixels exceeding the threshold were classified as foreground.

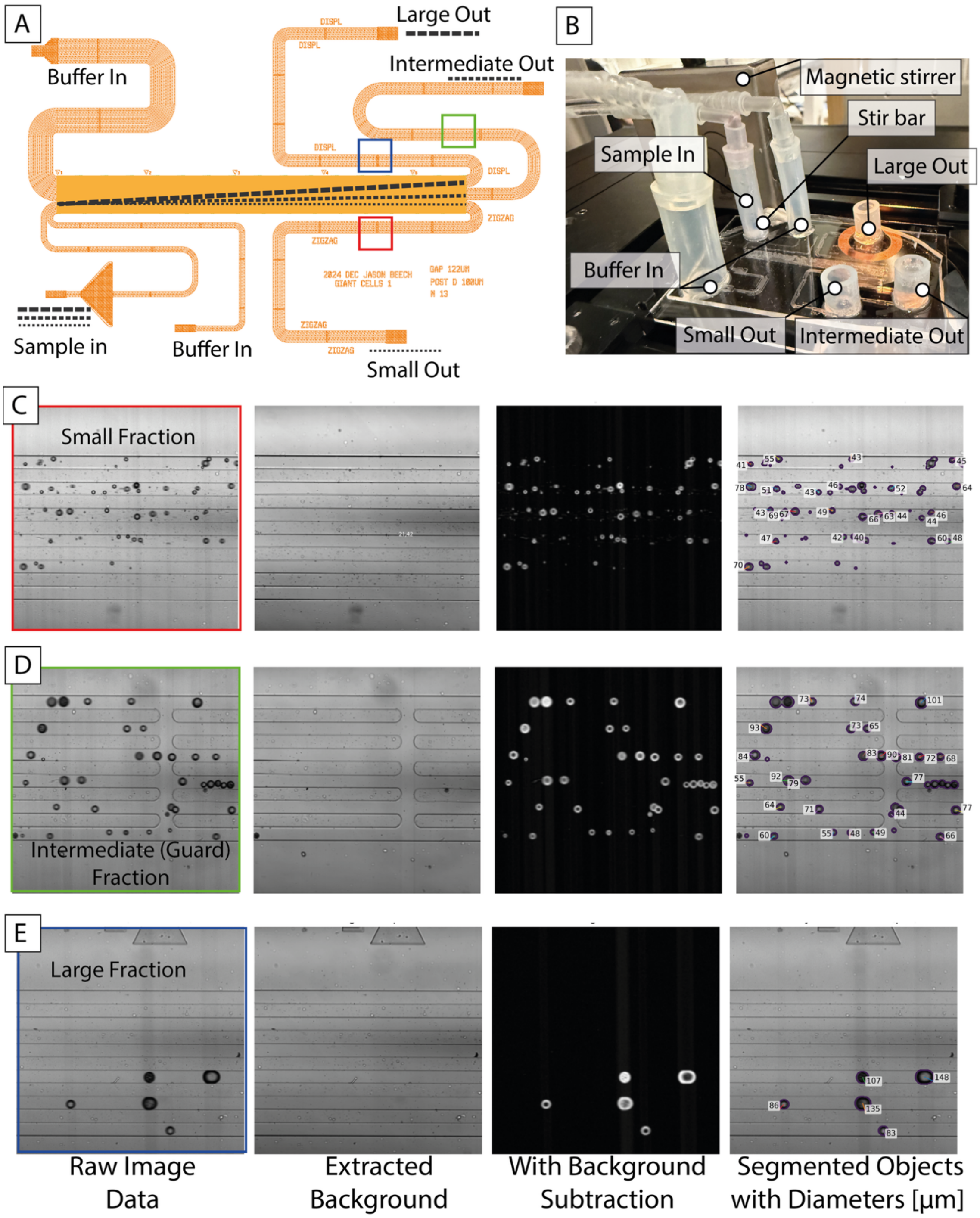


***Figure S1*** *Overview of device, image acquisition and image analysis. (A) Schematic of device showing how the sample is split into three size-based fractions in three outlets. (B) Photograph of the actual device where sample inlets/outlets can be seen. A small magnetic stir bar is used in the sample inlet reservoir to stop the fat cells separating from the buffer due to the large difference in bouyancy. (C, D, E) Data acquisition and analysis. Images aquired in the outlet channels, shown by red, blue, green squares in (A) furthest left. Moving right, calculated backgrounds, images of fat cells with background removed, and segmented and identified fat cells with diameters marked.*

#### Morphological cleanup

Binary masks were refined using standard morphological operations: a single iteration of binary opening to remove isolated noise pixels, followed by binary closing to fill small gaps, and hole filling to ensure contiguous particle regions. This produced clean binary masks corresponding to candidate particle objects.

#### Size-based pre-filtering

Connected components were labeled using 8-connectivity. To reduce false detections and accelerate downstream analysis, objects were filtered by area using a strong prior on expected particle size. Only objects with equivalent diameters between 5 and 120 pixels were retained. The equivalent diameter was defined as the diameter of a circle with the same area as the segmented object.

#### Shape measurement

For each retained object, geometric features were computed directly from the binary mask. Object area was measured as the number of foreground pixels. The perimeter was estimated from the difference between the object mask and its one-pixel erosion. From these quantities, circularity was calculated as

$$C = \frac{4\pi A}{P^2},$$

where $A$ is the object area and $P$ is the perimeter. Circularity approaches unity for ideal circular objects and decreases for elongated or merged shapes.

To further quantify object elongation, a moment-based eccentricity proxy was computed from the second central moments of the object mask. The eigenvalues of the corresponding covariance matrix were used to define an eccentricity-like metric ranging from 0 (circular) to 1 (highly elongated). This metric is robust to pixelation and does not require explicit contour fitting.

#### Decoupled filtering and analysis

Importantly, segmentation and geometric measurement were performed once per dataset, and all per-particle features were stored in a tabular format. Shape-based filtering (e.g., circularity or eccentricity thresholds) and downstream statistical analyses were applied in separate steps, enabling rapid iteration over filtering parameters without re-running the computationally expensive segmentation.

#### Particle Filtering and Preprocessing

Raw particle data were exported as a CSV file containing equivalent diameter ($D_{\text{eq}}$, μm), circularity, eccentricity proxy, and outlet label. All analyses were performed in Python using NumPy, Pandas, SciPy, and Matplotlib.

Particles were filtered prior to statistical analysis using both size and shape criteria:

1. **Diameter filtering:**
   Particles were restricted to

$$15\ \mu\text{m} < D_{\text{eq}} < 140\ \mu\text{m}$$

   to remove segmentation artifacts and large aggregates outside the physically relevant size range i.e. larger than smallest expected cell size and small enough to squeeze through the microfluidics channels.

2. **Shape filtering (sphericity selection):**
   To exclude non-spherical or poorly segmented objects, particles were required to satisfy:
   - Circularity ≥ 0.85
   - Eccentricity proxy ≤ 0.5

   Circularity is defined as $4\pi A/P^2$, where $A$ is particle area and $P$ is perimeter. A perfect circle has circularity = 1. These thresholds preferentially retain near-spherical particles while removing elongated or irregular objects.

#### Routing Probability Estimation

Routing behavior was quantified by estimating the conditional probability

$$P(\text{outlet} \mid D)$$

for each outlet as a function of particle diameter.

Diameter space was discretized into 40 equally spaced bins spanning the filtered diameter range. For each bin, the probability was computed as:

$$P_i(D) = \frac{n_i(D) + \alpha}{\sum_j n_j\,(D) + \alpha N}$$

where:

- $n_i(D)$ is the particle count in outlet $i$ within the bin,
- $N = 3$ is the number of outlets,
- $\alpha = 0.5$ is a Laplace smoothing parameter to avoid zero-probability artifacts.

Probability curves were smoothed along the diameter axis using a Gaussian filter ($\sigma = 1.2$ bins). After smoothing, probabilities were renormalized such that the sum across outlets equaled unity at each diameter.

#### Determination of the Cutoff Diameter $D_c$

The cutoff diameter $D_c$ separating the Small and Large outlets was defined as the diameter at which:

$$P(\text{Small} \mid D_c) = P(\text{Large} \mid D_c)$$

The intersection point was identified via sign change detection in the difference between the two probability curves, followed by linear interpolation between adjacent bins.

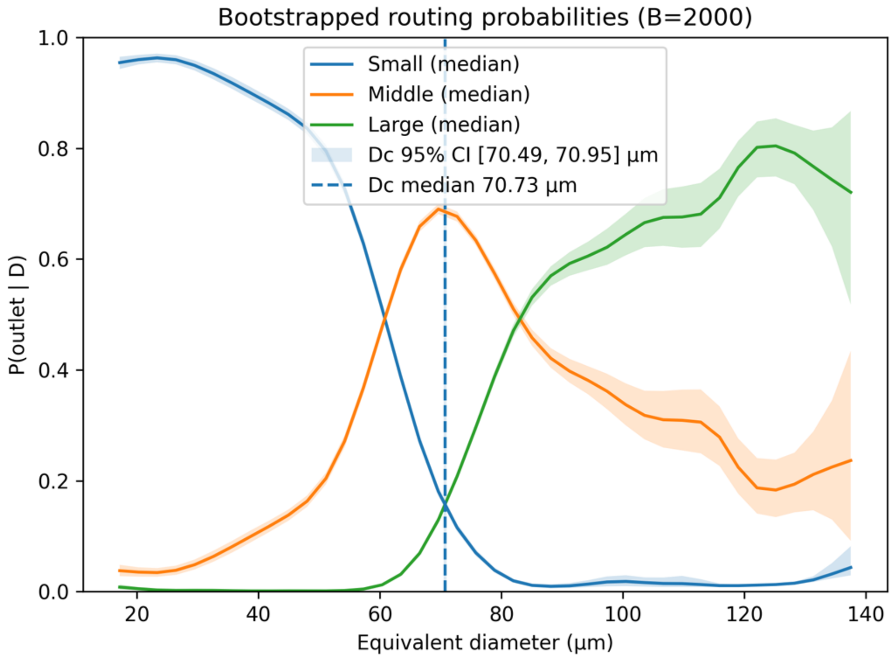


***Figure S2*** *Bootstrapped routing probabilities as a function of equivalent particle diameter. Solid lines represent the median conditional probability $P(outlet \mid D)$ across 2000 bootstrap replicates. Shaded regions indicate 95% confidence intervals. The vertical dashed line denotes the median cutoff diameter $D_c$ separating Small and Large outlets, and the shaded vertical band represents its 95% confidence interval.*

### Bootstrap Uncertainty Estimation

Uncertainty in routing probabilities and $D_c$ was quantified via stratified bootstrap resampling.

For each bootstrap replicate (B = 2000):

- Particle diameters were resampled with replacement independently within each outlet.
- Routing probabilities were recalculated.
- A new $D_c$ value was computed.

This stratified approach preserves the original outlet population sizes while estimating sampling variability.

From the bootstrap ensemble, the following were computed:

- Median probability curves
- 95% confidence intervals (2.5–97.5 percentiles)
- Median and 95% confidence interval of $D_c$

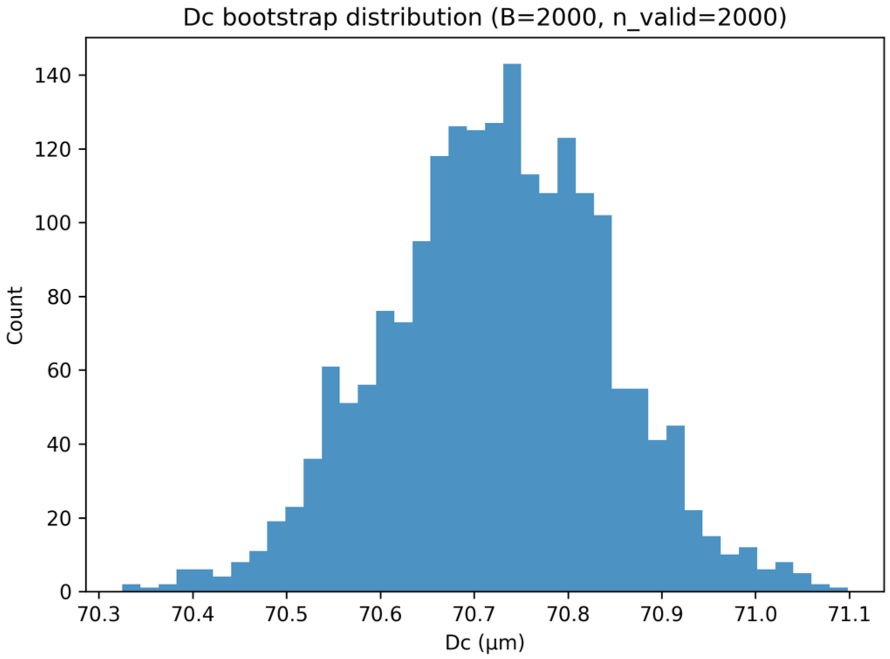


***Figure S3*** *Bootstrap distribution of the cutoff diameter $D_c$ (Small–Large crossover). The histogram represents 500 stratified bootstrap replicates. The distribution quantifies sampling uncertainty in the inferred size threshold.*

### Purity and Recovery Metrics

Using the bootstrap-derived $D_c$, classification performance for the Small and Large outlets was evaluated.

Particles were classified as:

- Small if $D < D_c$
- Large if $D \geq D_c$

For each outlet:

- **Purity** was defined as the fraction of particles collected in that outlet that were correctly classified by size.
- **Recovery** was defined as the fraction of particles predicted to belong to that size class that originated from the corresponding outlet.

Bootstrap distributions of purity and recovery were used to compute means and 95% confidence intervals.

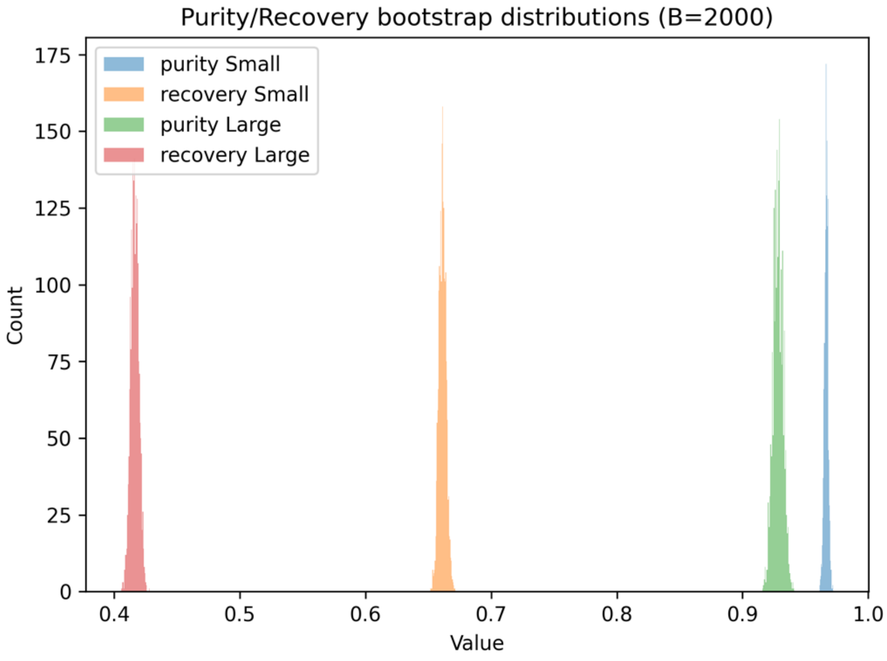


***Figure S4*** *Bootstrap distributions of purity and recovery for Small and Large outlets based on the size threshold $D_c$. Histograms represent 2000 bootstrap replicates. Median values and 95% confidence intervals are reported in Table S1.*

***Table S1*** *Summary statistics for the cutoff diameter $D_c$ and classification performance metrics derived from stratified bootstrap analysis (B = 2000). Reported values include the mean, standard deviation, median (p50), and 95% confidence interval (2.5–97.5 percentiles). Purity and recovery are defined with respect to the diameter-based classification using the bootstrap-derived $D_c$.*

| | mean | std | P 2.5 | P 50 | P 97.5 | N |
|---|---|---|---|---|---|---|
| Dc [µm] | 70.725 | 0.119 | 70.491 | 70.726 | 70.955 | 2000 |
| Purity (Small) | 0.967 | 0.002 | 0.963 | 0.967 | 0.970 | 2000 |
| Recovery (Small) | 0.661 | 0.003 | 0.656 | 0.661 | 0.667 | 2000 |
| Purity (Large) | 0.928 | 0.004 | 0.921 | 0.928 | 0.936 | 2000 |
| Recovery (Large) | 0.417 | 0.003 | 0.410 | 0.416 | 0.423 | 2000 |

### Movies

Movies can be found here: https://doi.org/10.5446/s_2017

**Movie 1 – External view of DLD device size-sorting adipocytes**

The movie is taken with a cell phone and shows an external view of a DLD device in action size-sorting adipocytes.

**Movie 2 – Following adipocytes through a DLD device as they are size-separated**

This movie, taken through a microscope, follows adipocytes as they travel through the device becoming separated by size.

**Movie 3 – Imaging and automated size measurement of adipocytes, post size-sorting by DLD**

Adipocytes imaged after separation but before leaving the device. Video also shows examples of segmentation results from which diameters are extracted using a custom image analysis pipeline written in Python.

### Code

All code for image segmentation and the extraction of cell sizes can be found here: https://doi.org/10.5281/zenodo.18789356

### Data

All data can be found here: https://doi.org/10.7910/DVN/M6DMVF
